# A simple analytical formula to compute the residual Mutual Information between pairs of data vectors

**DOI:** 10.1101/041988

**Authors:** Jens Kleinjung, Ton C.C. Coolen

**Affiliations:** The Francis Crick Institute, Mill Hill Laboratory, London NW7 1AA, U.K.; Institute for Mathematical and Molecular Biomedicine, Hodgkin Building, room HB 4.5N, Guy’s Campus, London SE1 1UL, U.K. and Department of Mathematics, Strand Building, room S5.26, Strand Campus, London WC2R 2LS, U.K.

## Abstract

**Summary:** The Mutual Information of pairs of data vectors, for example sequence alignment positions or gene expression profiles, is a quantitative measure of the interdependence between the data. However, data vectors based on a finite number of samples retain non-zero Mutual Information values even for completely random data, which is referred to as background or residual Mutual Information. Estimates of the residual Mutual Information have so far been obtained through heuristic or numerical approximations. Here we introduce a simple analytical formula for the computation of the residual Mutual Information that yields precise values and does not require the joint probabilities between the vector elements as input.

**Availability and Implementation:** A C program *arMI* is available at http://mathbio.crick.ac.uk/wiki/Software#arMI. Using an input alignment in FASTA format or alternatively an internally created random alignment of specified length and depth, the program computes three types of Mutual information: *(i)* Shannon’s Mutual Information between all pairs of alignment columns; *(ii)* the numerical residual Mutual Information by using the same formula on the randomised (shuffled) data; *(iii)* the analytical residual Mutual Information introduced here. The package depends on the GNU Scientific Library, which is used for vector and matrix operations, factorial expressions and random number generation (Galassi *et al.*, 2009). Reference alignments and result data are included in the program package in the folder ‘tests’. The R environment was used for statistics and plotting (R Core Team, 2014).

**Supplementary Material:** A detailed derivation of the analytical formula is given in the Supplementary Material.

---

Structural and functional constraints have shaped biomolecules and their functions over evolutionary times. This is reflected for example in positional conservation of nucleotide and amino acid sequences across multiple species or in gene expression profiles present in related cell states. These types of correlation patterns can be detected by computing the Mutual Information (*I*_*x, y*_) between pairs of data vectors. Shannon’s Mutual Information *I*_*x, y*_ = 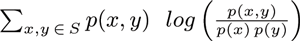 (Cover and Thomas, 2006) scales the joint probability *p*(*x, y*) to observe a specific element pair *x* and *y* with the marginal probabilities *p*(*x*) and *p*(*y*). *S* is a base set of symbols (alphabet) onto which the vector elements are mapped; for biological sequence alignments that corresponds generally to 4 nucleotides (DNA, RNA) or 20 amino acids (proteins) and for expression profiles typically to the number of genes under study. We will use *I*_*x, y*_ is this sense throughout the paper.

A critical question for all methods using correlation signals is whether the signal strength of the observed *I*_*x, y*_ is above the background or residual Mutual Information 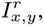 which is the Mutual Information between two fully independent alignment positions. The theoretically desirable 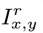 value of zero is obtained with a hypothetical random sample of infinite size, while in real biological data two error sources lead to non-zero background levels: i) under-sampling due to finite sample size and ii) redundancy among data due to their phylogenetic or functional relatedness, both yielding sampled frequency probabilities 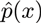, 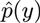 and 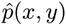 differing from expectation values. Heuristic methods have been developed to estimate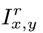 or to derive covariation values that have been corrected for background signals. The ‘average product correlation’ evaluates a form of excess *I*_*x, y*_ of two alignment positions *versus* the average *I*_*x, y*_ over all pairs of alignment positions (Dunn *et al.*, 2008). Alternatively, the covariance of alignment positions can be quantified *via* estimation of the sparse inverse covariance (Meinshausen and Bühlmann, 2006). A term for the expected systematic error of *I*_*x, y*_ has been proposed by Roulston (1999). A numerically inspired estimator of 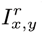 is the *I*_*x, y*_ value obtained from randomised (shuffled) data (Hempel *et al.*, 2011). In the following we will use this numerical residual Mutual Information 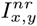 for comparison.

Here we present a simple analytical formula to compute the analytical residual Mutual Information 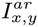 that has been derived from Shannon’s formula under the basic assumption that 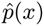 and 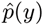 were statistically independent of each other, which is the essential condition to obtain 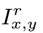 values (see Supplementary Material for the detailed derivation):

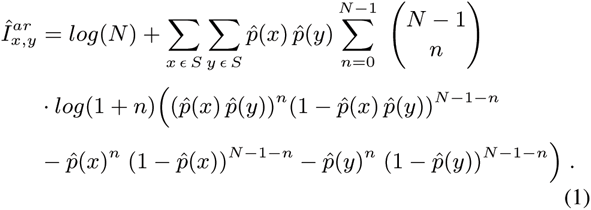

Equation 1 depends only on the sampled element probabilities 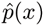 and 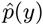 and on the vector length (or sample size) N. The assumption of statistical independence has led to the elimination of the joint probabilities *p*(*x, y*) occurring in Shannon’s *I*_*x, y*_ formula, simplifying the input to the probabilities of the base set symbols. This has favourable practical implications, for example in sequence analysis and design, where 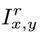 can now be controlled by variation of the composition of alignment profiles without the need to actually create these alignments.

A direct comparison between the 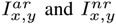 measure, unbiased by input data correlation, was obtained from a random sequence alignment of length 10 and depths (sample sizes) between 20 and 1000. Figure 1a shows the root mean square deviation between 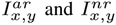 over all computed values. It is apparent that the numerical estimator 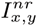 significantly overestimates 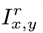 below sample sizes of about 500. However, the computational time per pair of data vectors is almost unaffected by the sample size for 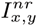 (Figure 1b), while it increases linearly with the sample size for the analytical estimator 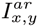. The absolute value of 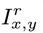 depends on the distribution of the element probabilities in the base set as shown in Figure 1c, where the total probability is evenly spread over varying base set sizes from 1 to 20 elements (sample size 100). The absolute 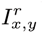 values and also the difference between 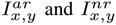 increase with the size of the base set.

To illustrate the application of 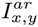 (equation 1) on biological data, a short and gap-less alignment of the switch II region (residues 57-66) of the Ras protein was taken from the Pfam database (Finn *et al.*, 2014). The chosen sample size was 100 to emphasise the differences between 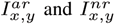 We computed the three described types of *I*_*x, y*_ between all column pairs: *(i)* Shannon’s *I*_*x, y*_ of the original alignments, yielding the biological correlation signal (uncorrected for 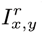); (ii)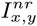 by application of Shannon’s *I*_*x, y*_ formula on shuffled alignment columns, where randomisations were repeated 100 times to estimate the mean and variance; *(iii)*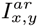 computed according to equation 1. Figure 1d shows the resulting *I*_*x, y*_ (black diamonds), 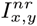 (orange circles, mean±sd) and 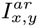(blue triangles) values plotted over the array of all column combinations (1:2, 1:3,…,1:10, 2:3, 2:4,…,9:10) for sample size 100. *I*_*x, y*_ values fluctuate depending on the correlation between the particular combinations of alignment columns. arMI yields the analytically correct residual values, while 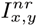 overestimates the background correlation as described above.

**Fig. 1.**
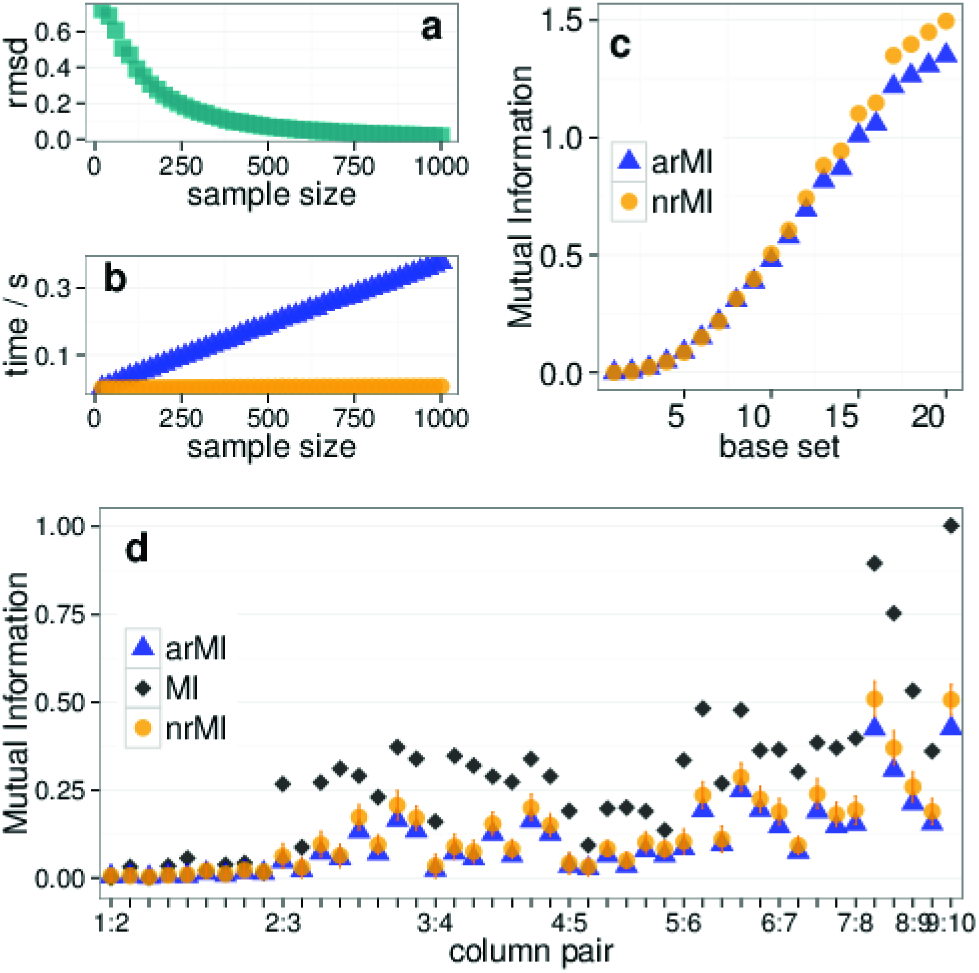
(a) Root mean square deviation between 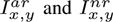 values for random sequence alignments of length 10 and depth (sample size) 20 to 1000 and (b) the computational time per vector pair (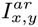: blue triangles, 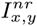: orange circles). (c) 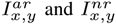 base set sizes 1 to 20 with even element probabilities. (d) Comparison between 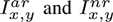 values of an alignment of 100 sequences of the protein Ras (residues 57-66).

In conclusion, the analytical 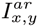 equation 1 is a precise and practical estimator of the residual *I*_*x, y*_ in particular for sample sizes below 500, where the usually employed numerical randomisation deviates from appreciably from the expectation values. The results suggest a pragmatic strategy for the computation of 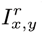, which is to use the analytical formula for smaller samples and the numerical approach for larger samples, because that strategy should yield a high precision of the resulting 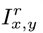 at low computational costs across a wide range of sample sizes. Due to the fact that the joint probabilities *p*(*x, y*) have been eliminated from the analytical equation, the 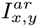 measure provides a means to explore 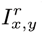 by varying vector compositions without explicitly pairing the vector elements.

## Time Complexity

The time complextity of the underlying algorithms has two main components, the combinatorics of the column pair comparisons and the MI calculation. The former, *N* * (*N* − 1)/2 pair comparisons, have a time complexity of *O*(*N*^2^), which applies to both, 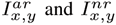 computations.

Disregarding the combinatorial part, we focus on the MI computation of single column pairs. The MI computation is dominated by random shuffling in the case of 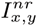 and by the evaluation of polynomial terms in the case of 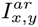.

Random shuffling was performed using the GSL function ‘gsl_ran_shuffle’, which is an implementation of the Durstenfeld version of the FisherYates shuffle. The algorithm has time complexity O(N), where *N* is the number of elements in the set (Durstenfeld, 420) or the sample size in our context.

Contrastingly, the computational time spent on the 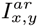 computation is dominated by the polynomials *p(x)p(y)*^*n*^ and *p*(*x*)*p*(*y*)^*N*−1−*n*^, which are evaluated (*N* − 1) times. Therefore, the time complexity is also O(N), but the actual time spent on the respective subroutines is considerably different, with random shuffling being about 25 times faster than evaluation of polynomial terms at large N (Fig. 2).

**Fig. 2.**
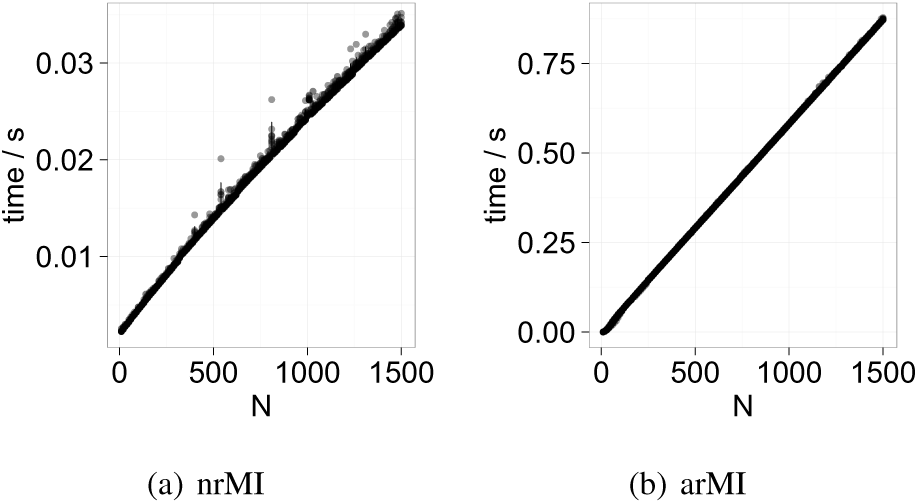
Computational time for the computation of (a) 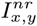 and (b)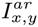 for a single column pair of sample size *N*. Each computation is repeated 10 times. The standard deviation is indicated by vertical bars.

## ROC curve

To evaluate the difference in performance between 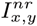 and 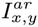 benchmark test on biological data was performed.

## ACKNOWLEDGEMENT

*Funding*: JK acknowledges support by the Francis Crick Institute (U117581331).

